# Improved data sets and evaluation methods for the automatic prediction of DNA-binding proteins

**DOI:** 10.1101/2021.04.09.439184

**Authors:** Alexander Zaitzeff, Nicholas Leiby, Francis C. Motta, Steven B. Haase, Jedediah M. Singer

**Affiliations:** Two Six Technologies, Arlington, Virginia, United States of America; Department of Mathematical Sciences, Florida Atlantic University, Boca Raton, Florida, United States of America; Department of Biology, Duke University, Durham, North Carolina, United States of America

## Abstract

**Motivation:** Accurate automatic annotation of protein function relies on both innovative models and robust datasets. Due to their importance in biological processes, the identification of DNA-binding proteins directly from protein sequence has been the focus of many studies. However, the data sets used to train and evaluate these methods have suffered from substantial flaws. We describe some of the weaknesses of the data sets used in previous DNA-binding protein literature and provide several new data sets addressing these problems. We suggest new evaluative benchmark tasks that more realistically assess real-world performance for protein annotation models. We propose a simple new model for the prediction of DNA-binding proteins and compare its performance on the improved data sets to two previously published models. Additionally, we provide extensive tests showing how the best models predict across taxonomies.

**Results:** Our new gradient boosting model, which uses features derived from a published protein language model, outperforms the earlier models. Perhaps surprisingly, so does a baseline nearest neighbor model using BLAST percent identity. We evaluate the sensitivity of these models to perturbations of DNA-binding regions and control regions of protein sequences. The successful data-driven models learn to focus on DNA-binding regions. When predicting across taxonomies, the best models are highly accurate across species in the same kingdom and can provide some information when predicting across kingdoms.

**Code and Data Availability:** All the code and data for this paper can be found at https://github.com/AZaitzeff/tools_for_dna_binding_proteins.

## 1 Introduction

There are over 195 million known natural protein sequences that have not been experimentally annotated for function (Consortium, 2018). Our ability to sequence genomes and identify proteins outstrips our ability to experimentally identify their functions. As such, there is a need for tools to predict protein function directly from sequence. In this paper, we focus on predicting whether a protein binds to DNA.

DNA-binding proteins play important roles in cellular processes including chromosomal DNA organization, initiation and regulation of transcription, DNA replication, DNA recombination, and DNA modification (Jones *et al*., 1987; Jen and Travers, 2013). Accurate identification of DNA-binding proteins has implications in both systems and synthetic biology by, for example, reducing the hypothesis space of potential control elements in gene regulatory networks, or by expanding the collection of known binders that could elucidate principles of design and thereby improve *de novo* design of DNA binders. Typically gene function is inferred from a combination of experimental, phylogenetic, and/or computational evidence (Giglio *et al*., 2019). While experimental evidence may be the most reliable, it is also the most costly and far slower than genome sequencing. Thus, computational methods that infer annotations directly from sequence, and thereby sub-select and prioritize specific targets for experimental validation, could increase our efficacy in identifying DNA-binding elements in novel organisms and designing new binders.

Recently there has been an explosion of models to predict DNA-binding proteins from sequence-based properties (Kumar *et al*., 2007; Lou *et al*., 2014; Xu *et al*., 2014; Liu *et al*., 2014, 2015; Motion *et al*., 2015; Liu et al., 2016; Waris *et al*., 2016; Wei *et al*., 2017; Qu *et al*., 2017; Zaman *et al*., 2017; Chowdhury *et al*., 2017; Zhang and Liu, 2017; Rahman *et al*., 2018; Du *et al*., 2019; Adilina *et al*., 2019; Ali *et al*., 2019; Mishra *et al*., 2019; Hu *et al*., 2019; Wang *et al*., 2020). Thus a fundamental question is how to assess the relative accuracy of proposed models predicting DNA-binding proteins. One reasonable method is to take experimentally labeled proteins, divide the sequences into a training and test set, and compare all the models on this train/test task. In current literature, the most common train/test task used to compare models is to train on the set of protein sequences in the “PDB1075” data set by (Liu *et al*., 2014, 2015) (hereafter referred to as the “benchmark training set”) and predict on the “186PDB” data set (Lou *et al*., 2014) (hereafter called the “benchmark test set”). The benchmark training set and benchmark test set data sets have been used extensively (Liu *et al*., 2016; Wei *et al*., 2017; Zaman *et al*., 2017; Chowdhury *et al*., 2017; Zhang and Liu, 2017; Wang *et al*., 2017; Rahman *et al*., 2018; Hu *et al*., 2019; Du *et al*., 2019; Adilina *et al*., 2019; Ali *et al*., 2019; Mishra *et al*., 2019; Wang *et al*., 2020), with each new model showing improvements on the task. Unfortunately, there are flaws with this train/test task:

- 75 of the93 (80.6%)DNA-binding protein sequences in the benchmark test set are also in the benchmark training set. While some papers (Liu *et al*., 2014, 2015; Zhang and Liu, 2017) remove sequences from the benchmark training set that are above a certain percent identity to those in the benchmark test set, it is unclear (and unlikely) that all papers do so. When a protein is present in both the training and the test set, models are rewarded for memorizing those proteins (often over-fitting) rather than learning general principles of DNA binding. Thus, the reported results may not accurately reflect the expected future performance if and when these models are applied to novel sequences not contained in the training or test sets.
- The benchmark training set contains duplicate entries for identical monomers making up homodimers and homotrimers: entries 2AY0A, 2AY0B, and 2AY0C are the same, as are 3FDQB and 3FDQA, as well as 4GNXL and 4GNXK. While this does not invalidate predictions on the benchmark test set, most of the mentioned papers (Liu *et al*., 2014, 2015; Ma *et al*., 2016; Liu *et al*., 2016; Wei *et al*., 2017; Zaman *et al*., 2017; Chowdhury *et al*., 2017; Zhang and Liu, 2017; Wang *et al*., 2017; Rahman *et al*., 2018; Adilina *et al*., 2019; Ali *et al*., 2019; Mishra *et al*., 2019; Wang *et al*., 2020) report the results of jackknife testing on the training set, where they hold out each sample in turn for testing after training on the rest of the samples in the benchmark training set. This results in the duplicated proteins being both memorizable and over-represented in the evaluation of the model.
- Some entries in the benchmark training set are not protein sequences but DNA sequences: specifically entries 4GNXL, 4GNXK, 1AOII, 3THWD, 4FCYC, 4JJNJ, and 4JJNI. These sequences were rejected from the PDB due to being DNA sequences rather than amino acid sequences, but they remain in the benchmark training set and thus might be treated as DNA-binding proteins.

Another important consideration is that the benchmark training set is very small compared to the amount of available data. Machine learning models generally perform better, and exhibit greater generalizability, when trained on more data. The train/test task should mimic situations in which users would employ the models; having an unrealistically small benchmark training set makes it unclear whether benchmark model evaluations apply to actual use.

Other studies have used different data sets to predict DNA-binding proteins, but they too are either small (fewer than 200 DNA binders in the training set (Waris *et al*., 2016; Xu *et al*., 2014)), suffer from overlap between their training and testing set (Qu *et al*., 2017; Hu *et al*., 2019), or are unavailable for download (Peled *et al*., 2016). The authors of (Qu *et al*., 2017) acknowledge the overlap and create a training set they claim has no overlap with the testing set, but they do not provide the no-overlap training set publicly.

One contribution of the work herein is the assembly and partition of larger, more reliable training and test sets. We also provide code and instructions for generation of similar data sets to address other annotation problems. These sets contain no overlap between the train and test sets, have no duplication of sequences within sets, contain only amino acid sequences, and are at least 10 times larger than the benchmark training and test set. These sets were split at random, to mimic the generation of train/test splits in the aforementioned papers. This evaluation task approximates a scenario in which many DNA binders are known, and models are tasked with identifying binders that previous efforts have missed. Using these larger training and test sets, we show that two models substantially outperform recently published results: an off-the-shelf gradient boosting model using features from a sequence embedding by a protein language model (Rives *et al*., 2019), and a simple nearest neighbor model using BLAST percent identity.

A fundamental question of interest to biologists is whether a model could identify DNA binding proteins in a newly sequenced species with no functional annotations. To approximate this use case with an evaluative task, we ask how well a model can predict DNA binding for proteins of a single species held out from the training data. We evaluate the gradient boosting and nearest neighbor models (the best performing in the previous random train/test task) in a variety of such settings. The results show that the gradient boosting model using the sequence embedding features gives comparable results to the nearest neighbor model. Because of the untapped potential for refinement and tuning of this model, it may be a good starting point for future work in protein function prediction.

Our analysis reveals some of the weaknesses in the current body of work for predicting DNA-binding proteins. We propose several improvements to data handling and model evaluation for the prediction of DNA binding and for protein function prediction in general.

## 2 System and methods

### 2.1 Data sets

We propose several new data sets for DNA binding prediction tasks that address the limitations of the existing benchmark data sets. Our new data sets are large, cleaned to ensure data quality, divided into non-overlapping training and test sets, and available at the provided GitHub account. Additionally, we provide tools and instructions for making machine learning ready data sets for protein function prediction more generally.

The data are taken from UniProt (Consortium, 2018). We take all bacterial and eukaryotic proteins in the Swiss-Prot subset with a sequence length between 40 and 5600 (filters: taxonomy:2 or taxonomy:2759, reviewed:yes, length:[40 TO 5600]). We set this upper limit because the longest DNA-binding protein (defined below) in UniProt is 5,588 amino acids long. We exclude sequences that appear in multiple entries which disagree about whether the protein is DNA-binding. These filters on the UniProt database result in 419,206 unique protein sequences (as of Dec 10th, 2020). To determine whether a protein is DNA-binding, we use GO code (Ashburner *et al*., 2000; Consortium, 2019) 3677 (name: “DNA-binding”, definition: “Any molecular function by which a gene product interacts selectively and non-covalently with DNA”) and any of its children through an is-a or part-of relationship.

We use a random split of the data to build members of a suite of training/testing sets. We use these sets to compare performance between different models. A random split makes few assumptions on the data or prediction goal, and reflects most of what has been previously published on this topic. The predictive task best approximated by these data splits is a scenario in which many DNA binders are known, and models are used to identify potential additional binders that previous efforts have missed. First, we subset the data to only allow sequences with the 20 canonical amino acids. Then, we randomly select ≈ 10% of the proteins to be a base test set and the remainder to be a base training set. We derive three separate training/testing sets from this base split:

- **Random, length-limited (RLL)**: The training and test proteins all have sequence length ≤ 1000. We compare to two models from the literature that were built for sequences of limited length. To test the models under conditions similar to those published, we use these restricted length data sets.
- **Random, similarity-limited (RSL)**: We remove from the training set sequences that are similar to any sequence in the test set. The test set is the base test set, and the training set comprises only sequences for which the top five alignments (as measured by bit score) with each sequence in the testing set have ≤ 50% identity (under default BLAST parameters using BLOSUM62) (Altschul *et al*., 1990).
- **Random, similarity- and length-limited (RS&LL)**: A subset of RSL, with all sequences above length 1000 removed.

Table 1 details the number of sequences in each of the training and testing sets.

**Table 1.**
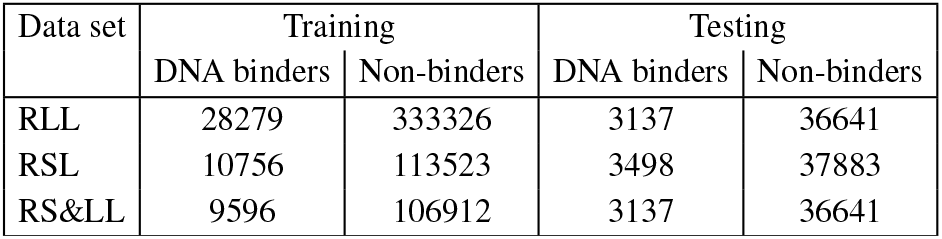
Number of sequences in the train and test sets.

We next make a suite of data sets geared towards the following use case: a new species has been sequenced and we want to use existing data to predict the molecular function of the proteins in this species. To mimic this situation, we choose the eight species in the bacterial kingdom and eight in the eukaryotic kingdom with the most annotated DNA binders. For each held-out species, we construct test sets of all amino acid sequences from that species, and three corresponding training sets: bacterial proteins, eukaryotic proteins, and bacterial and eukaryotic proteins. All sequences in the test set are excluded from all three training sets. Tables 2 and 3 enumerate each of the species we held out as well as the number of sequences that are DNA binders or non-binders in each one.

**Table 2.** Number of protein sequences for different bacterial species

| Species | DNA binders | Non-binders |
| --- | --- | --- |
| *Escherichia coli* | 996 | 8305 |
| *Salmonella enterica* | 490 | 5256 |
| *Staphylococcus aureus* | 332 | 2753 |
| *Mycobacterium tuberculosis* | 236 | 2195 |
| *Bacillus subtilis* | 424 | 3035 |
| *Streptococcus pyogenes* | 216 | 1595 |
| *Bacillus cereus* | 200 | 2419 |
| *Streptococcus pneumoniae* | 155 | 1451 |
| Total in kingdom | 18809 | 228415 |

**Table 3.** Number of sequences for different eukaryotic species

| Species | DNA binders | Non-binders |
| --- | --- | --- |
| *Homo sapiens* | 2448 | 16604 |
| *Arabidopsis thaliana* | 2018 | 13068 |
| *Mus musculus* | 1917 | 14614 |
| *Oryza sativa* | 765 | 3643 |
| *Rattus norvegicus* | 734 | 7217 |
| *Saccharomyces cerevisiae* | 641 | 5946 |
| *Xenopus laevis* | 570 | 2744 |
| *Drosophila melanogaster* | 480 | 3066 |
| Total in kingdom | 16448 | 155536 |

### 2.2 Metrics

In this section, we explain the metrics we use to compare models on the aforementioned data sets. Denote the number of true positives, true negatives, false positives, false negatives as TP, TN, FP, and FN respectively. Define the following:

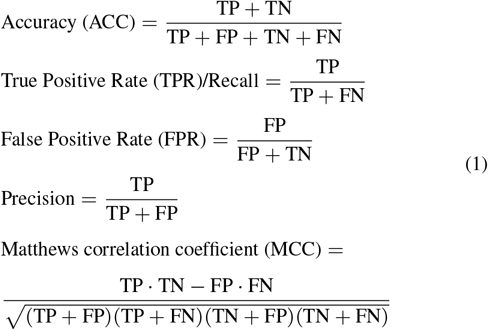

Matthews correlation coefficient (MCC) is a good measure when the data set is imbalanced (Chicco, 2017). However, it and the other metrics above require a binary positive or negative prediction, and the models we test predict probabilities. To use the above metrics, we have to choose a threshold such that a predicted probability at or above the threshold is a positive prediction (DNA-binding) and a predicted probability lower than the threshold is a negative prediction (non-DNA-binding). For all models in this paper, we take the threshold to be 0.5 when calculating ACC and MCC.

In addition, we include two metrics that evaluate the raw predicted probabilities: area under the receiver operating characteristic curve (ROC AUC) and average precision (AP). To define these, allow the quantities in equation (1) to be functions of the threshold *T*, e.g. TPR(*T*) is the true positive rate when threshold *T* is chosen. Then the definitions of ROC AUC and AP are

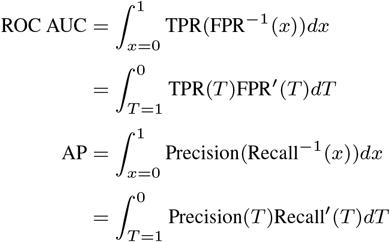

Like MCC, AP is a good metric for imbalanced data sets (Chicco, 2017).

## 3 Algorithms and Implementation

In this section, we describe the four different models that we compare.

As a baseline model, we use a nearest neighbor (1-NN) model based on BLAST percent identity (Altschul *et al*., 1990). The nearest neighbor model predicts each sequence in the testing set to have the same label as the sequence with the highest percent identity out of the five best alignments (as measured by bit score) with sequences in the training set. To calculate the percent identity and best alignments, we use BLAST software (Altschul *et al*., 1990) with default parameters using BLOSUM62. For more information about local alignment, see (Altschul and Gish, 1996).

Unfortunately, many of the previously mentioned papers do not have code available (Liu et al., 2014, 2015, 2016; Zaman *et al*., 2017; Wei *et al*., 2017; Chowdhury *et al*., 2017; Zhang and Liu, 2017; Xu *et al*., 2014; Peled *et al*., 2016), or have computationally expensive training procedures that would not scale well to a larger data set. For example, one of the methods in (Adilina *et al*., 2019) would require training on 32620 variants of the training set to determine the optimal features. As a result, we only evaluated two published modeling approaches, both neural networks: First, the model from (Qu *et al*., 2017), which features an embedding layer, two convolution layers, and an LSTM layer; second, the model from (Hu *et al*., 2019), which replaces the LSTM in (Qu *et al*., 2017) with 2 bi-LSTM layers.

In (Qu *et al*., 2017), the authors show the LSTM model performs better than (Kumar *et al*., 2007) and (Motion *et* al., 2015) on two test sets: one featuring 200 *Arabidopsis thaliana* proteins, the other featuring 200 *Saccharomyces cerevisiae* proteins. In (Hu *et al*., 2019), the authors give results of the bi-LSTM model and (Ma et al., 2016) on the aforementioned 200 *Arabidopsis thaliana* proteins, showing the bi-LSTM model performs the best. In addition, they show the bi-LSTM model has better results than (Motion *et al*., 2015; Ma *et al*., 2016; Qu *et al*., 2017) on a random test/train split. The bi-LSTM model is shown to have better performance on the benchmark test set than (Liu et al., 2014; Wang et al., 2017; Rahman *et al*., 2018; Chowdhury *et al*., 2017). It bears mentioning that it is unclear if the authors of (Qu *et al*., 2017; Hu *et al*., 2019) compare the models using the same training set in all of these results.

We train and make predictions using these models following the approaches of the authors (Hu *et al*., 2019; Qu *et al*., 2017). Specifically, for the LSTM model, we train 5 models using 5-fold cross-validation and use the predictions from the best performing model (as measured on the validation set). For the bi-LSTM model, we train three different models, one using 10% of the training data for validation, one using 15%, and the other using 20%. We use the predictions from the model that achieved the highest validation performance.

We also introduce a gradient boosting model trained with XGBoost (XGB) (Chen and Guestrin, 2016) applied to features generated from sequence data by the evolutionary scale model (ESM)(Rives *et al*., 2019). The ESM was trained on 250 million protein sequences in a self-supervised learning task to learn a language model for proteins. The ESM embeds each protein sequence as a 1280-feature vector purported to encode biochemical properties, biological variation, remote homology, and information in multiple sequence alignments. We use the features generated from ESM as-is, not tailoring them to our particular task.

For the gradient boosting model, we randomly sample 10% of the training data to serve as a validation set. The gradient boosting model sees the remainder of the training data at every step and we stop the training after the average precision fails to increase for ten rounds on the validation set; we then use the model from the best performing epoch. Positive samples in the training set are up-weighted so that positive and negative samples have equal aggregated weight. To account for the variance between individual runs, we train the model three times with different train/validation splits. The final prediction for each test sample is the median of the predictions of the three runs.

## 4 Results and Discussion

The nearest neighbor and gradient boosting models outperform the LSTM and bi-LSTM across all data sets. The nearest neighbor model, despite its simplicity, performs essentially as well as the gradient boosting model. We also investigate the sensitivity of each model to perturbations of DNA-binding and non-DNA-binding regions and find several different patterns.

Figures 1 to 3 contain the results on the three tasks from the random train/test split. Since the nearest neighbor model does not output probabilities, we only show its ACC and MCC score.

**Fig. 1:**
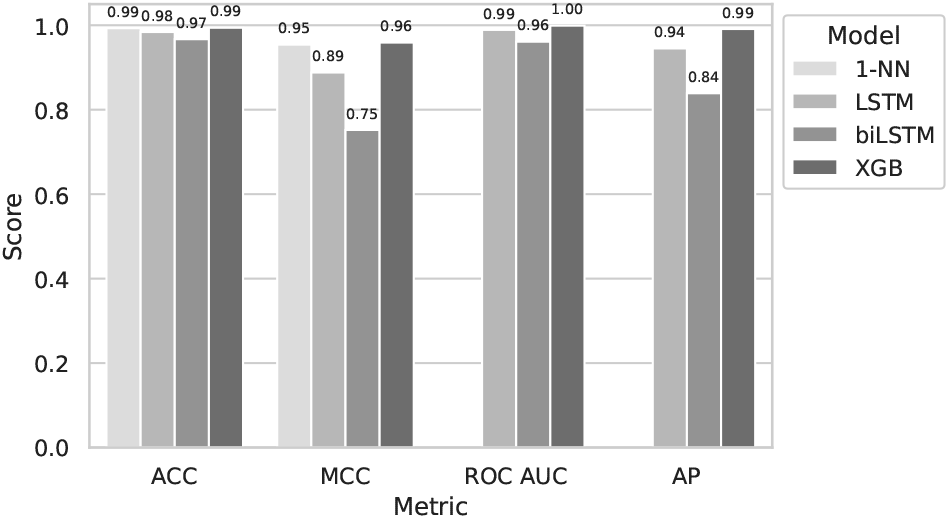
Performance on length-limited proteins (RLL split)

**Fig. 2:**
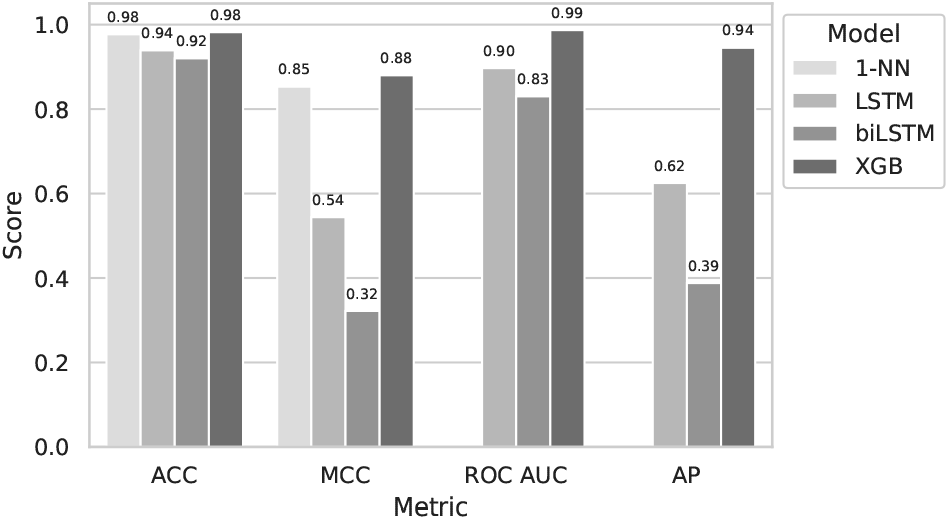
Performance on proteins with low train/test similarity (RSL split)

**Fig. 3:**
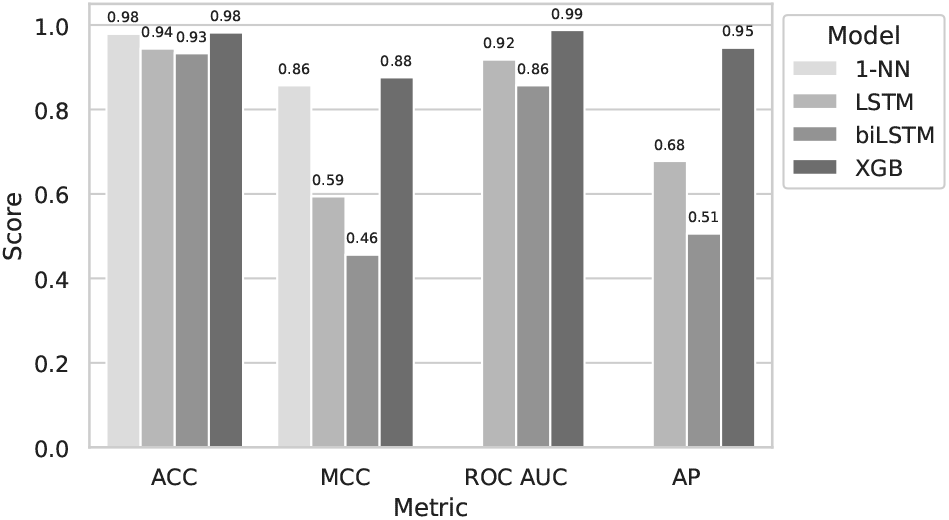
Performance on length-limited proteins with low train/test similarity (RS&LL split)

### 4.1 bi-LSTM performs worse than simpler LSTM

Notably, bi-LSTM performs worse than LSTM on all data sets. These results contrast with (Hu *et al*., 2019), where the authors present two train/test splits on which bi-LSTM has higher accuracy than LSTM. However, their first data set, which is similar to RLL, has about a 17% overlap between the training set and testing set. The second data set, which we could not obtain, has a small testing set and may contain overlap; the testing set is 200 sequences from the species *Arabidopsis thaliana* and their training set also contains sequences from *Arabidopsis thaliana*. Furthermore, it is not specified how the authors of (Hu *et al*., 2019) trained the LSTM model when comparing it with their bi-LSTM model. Different training approaches can yield models that perform differently. Performance differences may also arise because the training and test sets of (Hu *et al*., 2019) have a higher percentage of DNA-binding proteins (≈ 25%) than ours (≈ 8%) and are smaller (they are training with 57,078 total proteins vs our 361, 605 in RLL).

A weakness of the LSTM and bi-LSTM models is the fixed limit on sequence length that was imposed to keep the models from being too large. Figure 4 compares the performance of versions of the models with input size limits expanded so they could be trained on RSL (which is not size-limited) vs. RS&LL (which is size-limited) when predicting on the RS&LL test set (which is a subset of the RSL test set). We see that raising the maximum sequence length and expanding the training set of LSTM and bi-LSTM degrades the performance even on amino acid sequences of length ≤ 1000.

**Fig. 4:**
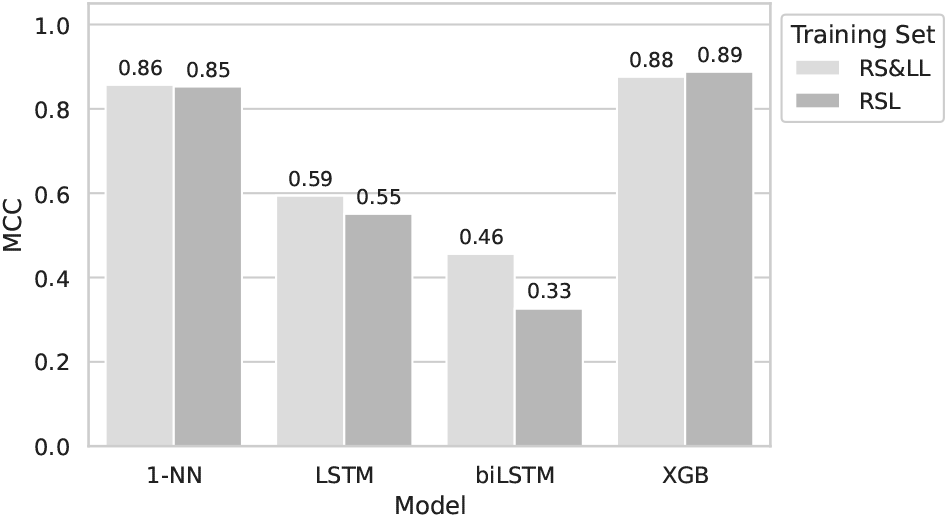
Mathews correlation coefficient on the RS&LL Test Set after training on RS&LL or RSL Training Set

### 4.2 Simple models outperform deep learning models

Unexpectedly, the BLAST nearest neighbor model does remarkably well on all the tasks; ACC and MCC are higher than both of the neural networks across all data sets, and often are competitive with the gradient boosting model (figures 1 to 3). One possible reason for the high performance of the nearest neighbor model is the way annotations in UniProt are generated. While for some proteins annotations reflect experimental observations, a search of Uniprot reveals that more than 97% of the DNA-binding data in the bacteria and eukaryotic kingdoms from Swiss-Prot are labeled by electronic annotation, biological aspect of ancestor, sequence or structural similarity, or sequence orthology. This could artificially boost performance: proteins are assigned a DNA-binding label due to sequence similarity, and then the nearest neighbor model predicts the same label also due to sequence similarity. To explore this possibility, we evaluated all four models on a subset of the RLL test set that included only positive examples with verified DNA binding activity (UniProt manual assertion codes EXP, IC, IDA, IPI, TAS, HDA) (figure 5). MCC dropped for all four models. The drop was particularly pronounced for the LSTM model, for which MCC decreased 25%. For the other three models, MCC decreased by 10%–14%. This test subset put bi-LSTM on par with LSTM, but the nearest neighbor and gradient boosted models remained substantially better. This suggests that the nearest neighbor model performance is not entirely explained by the electronic annotation process.

**Fig. 5:**
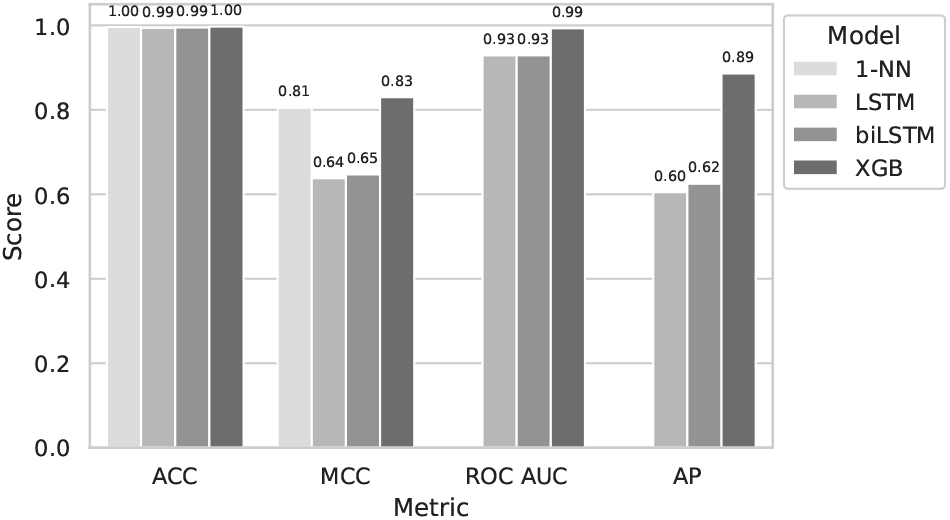
Performance on subset of length-limited proteins (RLL) where included positive examples have verified DNA binding activity

### 4.3 Model sensitivity to manipulation of DNA-binding regions

We next ask whether the models are identifying the DNA-binding regions of the proteins, as opposed to common patterns in DNA-binding proteins not specifically linked to DNA binding (e.g., nuclear localization sequences in eukaryotes (Görlich, 1997) and analogs in prokaryotes (Lisitsyna *et al*., 2020)). In the RS&LL test set, we examine 714 proteins that have a single DNA-binding region annotated. We modify their amino acid (AA) sequences in three ways to differentiate between these hypotheses:

- DBR reversed: we reverse the DNA-binding region (DBR) subsequence and leave the rest of the AA sequence intact.
- Random region reversed: we reverse a random subsequence of the same length as the DNA-binding region that does not overlap with the DNA-binding region. This test is a control for the effect of the reversal perturbation in general on model predictions.
- Everything but DBR reversed: we reverse the subsequences on either side of the DNA-binding region. This test determines the relative importance of the DNA-binding region compared to the rest of the protein sequence in determining model predictions.

We then compare models’ predictions on the modified amino acid sequences to those for the original protein sequences. We might expect that nearest neighbor model would not be greatly affected by reversing the DNA-binding region or a random region because the reversed region is a small portion of the entire sequence. In contrast, we expect that if the LSTM, bi-LSTM, and gradient boosting model have learned the importance of the DNA binding region in particular as opposed to their general context in the overall sequence, they would be more affected by reversing the DNA-binding region than by reversing a random region. By reversing both regions outside the DNA binding region we would expect the nearest neighbor model predictions to change dramatically, as much of the sequence has changed, but the impact on the other models would depend on the extent to which they had learned to identify DNA binding regions versus other patterns present in DNA binding proteins.

The results are shown in figure 6, where we plot the fraction of perturbed binders that each model classifies as binders. We observe three distinct patterns of response. The nearest neighbor and LSTM models show virtually no sensitivity to reversing a random region, modest sensitivity to reversing the entirety of the non-binding regions, and substantial sensitivity to reversing the DNA-binding region (figures 6a and 6b). This is consistent with the DNA-binding region being more highly conserved across proteins, with the BLAST alignment reflecting this conservation and the LSTM model learning to take advantage of it. The bi-LSTM model is moderately sensitive to reversing the entirety of the non-binding regions, and barely reacts to smaller perturbations to a random region or even the DNA binding region (figure 6c). This failure to react when the binding domain is disrupted is consistent with its failure to solve the binding prediction task. The gradient boosting model is insensitive to reverses of random non-binding sub-regions, but quite sensitive to reversing the DBR or the entirety of the non-binding regions (figure 6d). As with the LSTM, this likely reflects that the gradient boosting model has learned to focus on the binding domain. The nearest neighbor, LSTM, and gradient boosting models are all able to predict reversed DNA-binding domains as less likely to bind to DNA. The gradient boosted model, perhaps because it is built from an embedding that purports to holistically encode many features of proteins, is more sensitive to large disruptions of a protein’s non-binding components.

**Fig. 6:**
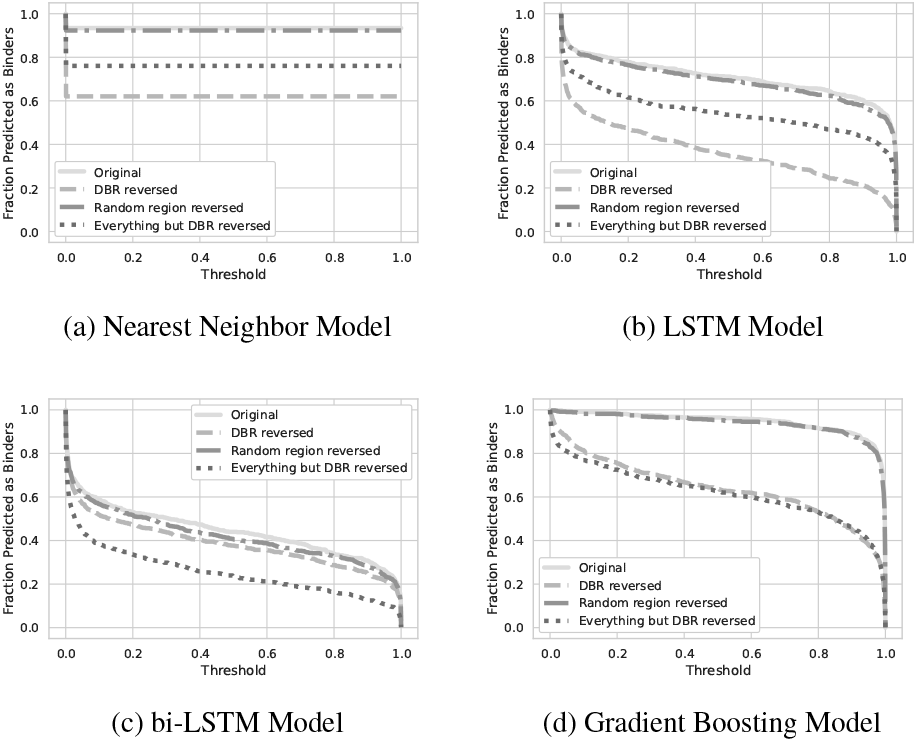
The effects of reversing different subsequences of DNA-binding proteins on the models.

### 4.4 Performance of nearest neighbor and gradient boosting models on data sets broken up by species

Next, we consider a set of tests that reflect the use case of annotating a novel proteome. Because the gradient boosting model performed the best on the previous sets and does not have a maximum length constraint, we use it in these tasks. Due to its simplicity and interpretability, we also consider the nearest neighbor baseline. We do not evaluate the LSTM or bi-LSTM models in these tasks, because their performance on the simpler tasks was substantially poorer and because the computational requirements to evaluate them on all of these tasks would have been quite large. First, we evaluate predictions for individual bacterial and eukaryotic species using training sets comprising each of the two kingdoms individually and both together. Figure 7 show the results for nearest neighbors and the gradient boosting model. Unsurprisingly, training on the “wrong” kingdom yields poorer results; however, we see evidence of information transfer across kingdoms in most cases. Training on both kingdoms yields results comparable to those obtained from training on only the kingdom of the test species. As with the previous results, the gradient boosting model and nearest neighbor model have comparable performance. Taken together with the previous results, the gradient boosting model with language features reaches an acceptable performance floor, indicating a good area for future research.

**Fig. 7:**
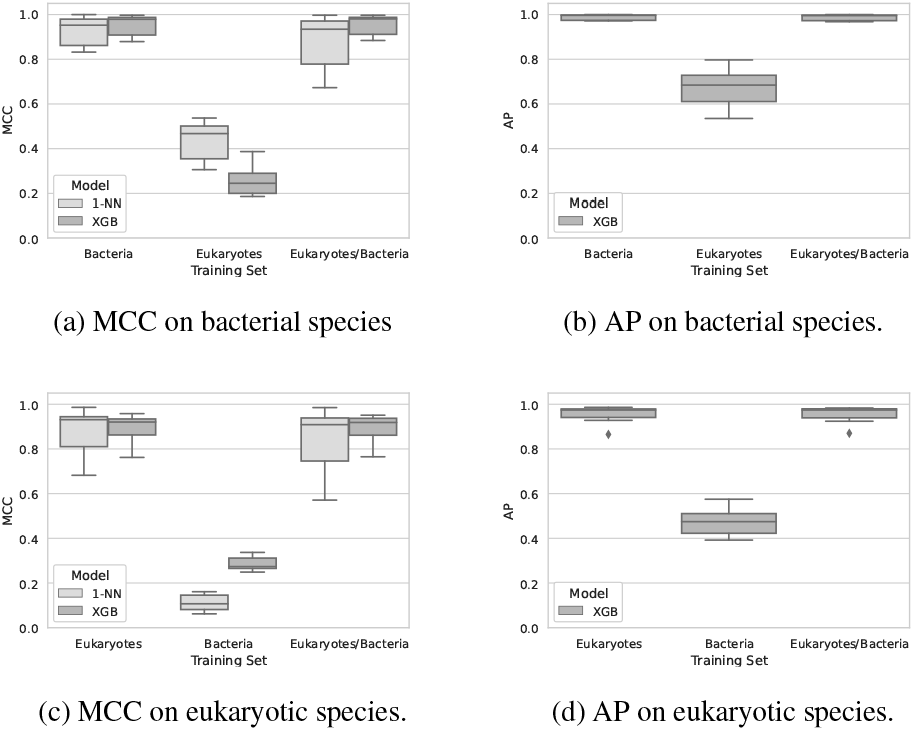
The Matthews correlation coefficient (MCC) for the nearest neighbor and gradient boosted models, and the average precision (AP) for the gradient boosted model, when predicting DNA-binding proteins for each of eight held-out test species per kingdom after training on data from bacteria, eukaryotes, and both together.

## 5 Conclusion

Automatic labeling of protein function is an important problem for computational biologists, as our ability to sequence genomes outstrips our ability to experimentally determine function. Focusing on predicting DNA-binding proteins, this paper makes four main contributions to the field.

First, we emphasize the importance of large, accessible data sets broken up into multiple training/testing tasks mirroring realistic use cases. Random train/test data splits, while common in the field of machine learning, don’t necessarily provide the best evaluation of predictive models. Data sets should be chosen with consideration for the scenarios in which a model is intended to be used. While having benchmark data sets is useful, as the underlying data compiled into these sets change, the benchmarks must be revisited. In the case of the PDB1075 test set widely used as a DNA-binding protein prediction benchmark, several data quality issues have never been corrected, and the data set remains relatively small. Here we offer a larger, carefully curated data set as a new standard that reflects the explosion of new sequence data at our disposal. We have also made available code for the straightforward construction of similar data sets.

Second, we showed that a simple baseline nearest neighbor model outperforms neural networks from recent literature in predicting DNA binding proteins. This held even when restricting training set sequences to those that were “far” from the test set, as well as when only using experimentally-labeled DNA binders to control for bias introduced by homology-based inferred labels in the UniProt data. While it may be surprising that BLAST percent identity outperforms more sophisticated modeling efforts, this is not inconsistent with our understanding of protein function. One paper notes that “several studies have shown that homologous sequences that share more than 40% identity are very likely to share functional similarity as judged by E.C. (Enzyme Commission) numbers” (Pearson, 2013). Because of its performance and simplicity, the BLAST nearest neighbor model predictions should serve as a baseline for future modeling efforts, and any proposed machine learning algorithm should be able to show a significant improvement over this baseline before it is considered as a replacement.

Third, the gradient boosted model we evaluated shows that a simple model based on a complex off-the-shelf embedding has acceptable performance on both the random split and held-out species data sets. While it does not perform substantially better than the nearest neighbor model, it shows promise as a foundation for a more capable model. We leave it to future work to determine if different model architectures can better utilize ESM features, or if other protein language model embeddings (e.g. ProtTrans (Elnaggar *et al*., 2020) or newer versions of ESM) yield better results.

Fourth, to better understand how the different models are making their predictions, we manipulated the amino acid sequences and measured the effect of these perturbations on the models. Our results indicate that the DNA-binding region is important to the predictions of the nearest neighbor, LSTM, and gradient boosting model. This suggests that the models are using information from the DNA binding regions of the proteins to make their predictions, as opposed to learning to recognize patterns in DNA-binding proteins apart from the binding domains.

As figure 1 and figure 7 show, for prediction of DNA-binding within kingdoms or within a random subset of the data, a simple single nearest neighbor BLAST and an out of the box gradient boosting model are near the performance ceiling. Practitioners can expect precision and recall of 94% for prediction in bacteria, and precision and recall of 90% for prediction in eukaryotes. Small tweaks to these simple models (e.g. majority vote of the five nearest neighbors) may push the prediction rate up slightly higher. There may be a place for advanced models in predicting DNA-binding proteins across kingdoms, but it is unclear how realistic these more difficult tasks are.

Future work should instead focus on the prediction of other protein functions where pattern identity methods (like BLAST) are inadequate (Ashkenazi *et al*., 2012). In these areas, more complex models have an opportunity to shine.

## Acknowledgments

Author roles by CRediT: Conceptualization: A.Z, F. M., J.S., N.L, and S. H. Data curation, software, investigation, visualization: A.Z. Writing – original draft, formal analysis, methodology: A.Z, J.S., N.L Writing – review & editing: All authors We are grateful for the use of computing resources and support from the Texas Advanced Computing Center (TACC).

## Funding

This work was supported by the Defense Advanced Research Projects Agency (DARPA) and the Air Force Research Laboratory under Contract No. [FA8750-17-C-0231] (and related contracts by SD2 Publication Consortium Members). Any opinions, findings and conclusions or recommendations expressed in this material are those of the authors and do not necessarily reflect the views of the Defense Advanced Research Projects Agency (DARPA), the Department of Defense, or the United States Government.

## Notes

https://github.com/AZaitzeff/tools_for_dna_binding_proteins

### Competing Interest Statement

The authors have declared no competing interest.

